# Altered Fhod3 Expression Involved in Progressive High-Frequency Hearing Loss via Dysregulation of Actin Polymerization Stoichiometry in The Cuticular Plate

**DOI:** 10.1101/2023.07.20.549974

**Authors:** Ely Cheikh Boussaty, Yuzuru Ninoyu, Leo Andrade, Qingzhong Li, Ryu Takeya, Hideki Sumimoto, Takahiro Ohyama, Karl J Wahlin, Uri Manor, Rick A Friedman

## Abstract

Age-related hearing loss (ARHL) is a common sensory impairment with complex underlying mechanisms. In our previous study, we performed a meta-analysis of genome-wide association studies (GWAS) in mice and identified a novel locus on chromosome 18 associated with ARHL specifically linked to a 32 kHz tone burst stimulus. Consequently, we investigated the role of Formin Homology 2 Domain Containing 3 (Fhod3), a newly discovered candidate gene for ARHL based on the GWAS results. We observed Fhod3 expression in auditory hair cells (HCs) and primarily localized at the cuticular plate (CP). To understand the functional implications of Fhod3 in the cochlea, we generated Fhod3 overexpression mice (*Pax2-Cre^+/-^; Fhod3^Tg/+^*) (TG) and HC-specific conditional knockout mice (*Atoh1-Cre^+/-^; Fhod3^fl/fl^*) (KO). Audiological assessments in TG mice demonstrated progressive high-frequency hearing loss, characterized by predominant loss of outer hair cells, and a decreased phalloidin intensities of CP. Ultrastructural analysis revealed shortened stereocilia in the basal turn cochlea. Importantly, the hearing and HC phenotype in TG mice were replicated in KO mice. These findings indicate that Fhod3 plays a critical role in regulating actin dynamics in CP and stereocilia. Further investigation of Fhod3 related hearing impairment mechanisms may facilitate the development of therapeutic strategies for ARHL in humans.

## 1 Introduction

Age-related hearing loss (ARHL) is one of the most common causes of sensorineural hearing loss among elderly population. Numerous studies have identified candidate genes associated with ARHL, implicating cellular pathways such as oxidative stress response (Sugiura et al. 2010; Someya et al. 2010), inflammation (Rodríguez-de la Rosa et al. 2017; Menardo et al. 2012), and cellular senescence (Benkafadar et al. 2019; Li et al. 2022). However, with the polygenic nature (Fransen et al. 2015) and phenotypic heterogeneity of ARHL (Gates and Mills 2005), the underlying mechanisms have not been fully elucidated. Further investigation is necessary to facilitate targeted interventions and personalized treatment approaches for ARHL.

Genome-wide association studies (GWAS) have been conducted in humans to explore the genetic architecture of ARHL and have successfully identified loci associated with susceptibility to ARHL (Friedman et al. 2009; Van Laer et al. 2010). However, GWAS studies often explain only a fraction of the heritability of complex traits, which imply the presence of undiscovered genetic variants contributing to ARHL. This may be attributed to limitations in study design, sample size, or the influence of rare variants that are challenging to detect. To address these challenges, we performed GWAS on a cohort of approximately 100 extensively characterized inbred mouse strains, known as the hybrid mouse diversity panel (HMDP) (Ghazalpour et al. 2012). This robust resource enables the investigation of both genetic and environmental influences on intricate traits. Our study successfully identified distinct genetic and phenotypic differences, expanding upon the known Mendelian ARHL genes (Ahl) described to date (Lavinsky et al. 2015; Ohmen et al. 2014). The mouse model provides a valuable platform for studying the genetics of hearing loss, as the mouse and human ears are functionally and genetically homologous, allowing for the translation of findings to human populations (Ghazalpour et al. 2012; Flint and Eskin 2012; Rau et al. 2015). However, additional studies are required to fully understand how the identified variants influence the ARHL phenotype. The loss of cochlear HCs and their associated mechanoelectrical transduction function significantly contributes to the development of ARHL (Schuknecht and Gacek 1993). HCs, specialized sensory cells within the inner ear, possess stereocilia, tiny hair-like structures arranged in a staircase pattern. Each stereocilium consists of a bundle of actin filaments, tightly regulated in length, and anchored at their base, into a cuticular plate that consists of dense actin meshwork. Actin-regulatory proteins play a crucial role in maintaining the fine actin structure, and mutations in these proteins have been associated with various forms of hereditary hearing loss (Miyoshi et al 2021). However, the precise role of actin-regulatory proteins in ARHL remains unclear. Rho-GTPases, such as cell division cycle 42 (Cdc42) and RhoA, are well-known actin-regulatory proteins that play a pivotal role in regulating actin dynamics in HCs (Ueyama et al 2014; Anttonen et al. 2017). Formin proteins that function as downstream effectors of RhoA are involved in regulating multiple cellular actin cytoskeletal elements by nucleating and polymerizing straight actin filaments (Breitsprecher and Goode 2013). While the Diaphanousrelated formin family (DRF) members DIAPH1 (Lynch et al. 1997) and DIAPH3 (Sánchez-Martínez et al. 2017; Schoen et al. 2010) are known hereditary hearing loss-related proteins, the physiological role of other DRF proteins in the HC, such as FHOD, have yet to be fully established.

FHOD3, initially described in the regulation of sarcomere organization in cardiomyocytes, has been localized to thin actin filaments with abundant expression in both myocardium and skeletal muscle (Taniguchi et al. 2009). Depletion of FHOD3 leads to a reduction of filamentous actin and disruption of sarcomere organization in cardiac cells (Taniguchi et al. 2009). Studies have shown that FHOD3 is involved in actin polymerization and myofibril integrity (Kan-o et al. 2012). Interestingly, FHOD3 has been observed at the pointed ends of actin filaments, where depolymerization normally occurs, suggesting a potential role in annealing short actin filaments (Fujimoto et al. 2016).

In this study, we aimed to investigate the functional implications of Fhod3, a novel candidate gene for ARHL identified through meta-analysis GWAS in mice. We generated two mouse models: Fhod3 overexpression mice (*Fhod3^Tg/+^; Pax2-Cre^+/-^*) and HC-specific conditional knockout mice (*Fhod3^fl/fl^; Atoh1-Cre*^+/-^) to assess the effects fluctuation in FHOD3 levels on the functioning of inner ear with audiological assessments in the transgenic mice revealing a progressive high-frequency hearing loss phenotype, accompanied by the predominant loss of outer HCs and deterioration of the cuticular plate. Scanning electron microscopy analysis demonstrated shortened stereocilia, particularly in the mid-base and base regions of the cochlea. Furthermore, targeted deletion of Fhod3 replicated the progressive hearing loss and HC deterioration observed in the Fhod3 overexpression mutants. These findings suggest a critical role for Fhod3 in regulating actin dynamics in the cuticular plate and stereocilia of high frequency cochlear hair cells.

## 2. Results

### 2.1 Random effects meta-analysis GWAS identifies 5 loci for ARHL in mice

Mouse strains exhibit higher heritability for complex traits, and the genetic loci associated with these traits tend to exert stronger effects compared to humans. Motivated by these advantages, we performed the first GWAS of its kind in the mouse by combining several data sets in a metaanalysis to identify loci associated with ARHL (Ohmen et al. 2014). We identified five genomewide significant loci (*P*<1.0×10^−6^). One of these confirmed a previously identified locus (*Ahl8*) on distal chromosome 11 and greatly narrowed the candidate region. Another locus of interest appeared for the 32 kHz tone burst stimulus and localized to chromosome 18 with the peak SNP (position 24320393; *P*=2.32 x 10^-6^) appearing within a haplotype block between 24.25 and 25.65 Mb containing the formin-like gene *Fhod3* (Ohmen et al. 2014). Due to its role in actin polymerization/capping, we decided to delve into the potential role of *Fhod3* in high-frequency and progressive hearing loss.

### 2.2 Spatial expression patterns of Fhod3 mRNA and protein in the cochlea

To analyze the gene expression patterns of Fhod3 in the inner ear, we employed multiplexed error-robust fluorescence in situ hybridization (MERFISH) (Chen et al. 2015) in P5 cochlea. The application of single-molecule imaging allowed us to visualize Fhod3 expression in auditory HCs and SGNs of the cochlea with high resolution (Fig. 1A to C). Additionally, we conducted immunofluorescence staining in P5 cochleae to further confirm the localization of Fhod3. Our results revealed the expression of Fhod3 in both outer HCs (OHCs), inner HCs (IHCs), and in SGNs (Fig. 1C). We generated a conditional knock-out mouse model targeting Fhod3 specifically in HCs using Atoh1-cre and Fhod3-flox mice. In these knock-out mice (*Fhod3^fl/fl^; Atoh1-Cre*^+/-^), Fhod3 signals were absent in HCs but still observed in SGNs, verifying Fhod3 expression in the cochlea (Fig.1C). Furthermore, to gain insights into the subcellular localization of Fhod3 within HCs, we developed an Fhod3-HA tagged knock-in mouse model (Fhod3^HA/+^) in which a 3xHA sequence was inserted into the C-terminus of Fhod3 prior to the stop codon. This demonstrated that Fhod3 predominantly localized to the cuticular plate in HCs (Fig.1D and E). Supporting these findings, data obtained from gEAR.org (https://umgear.org/) corroborates the high enrichment of Fhod3 expression specifically in cochlear HCs.

**Fig.1:**
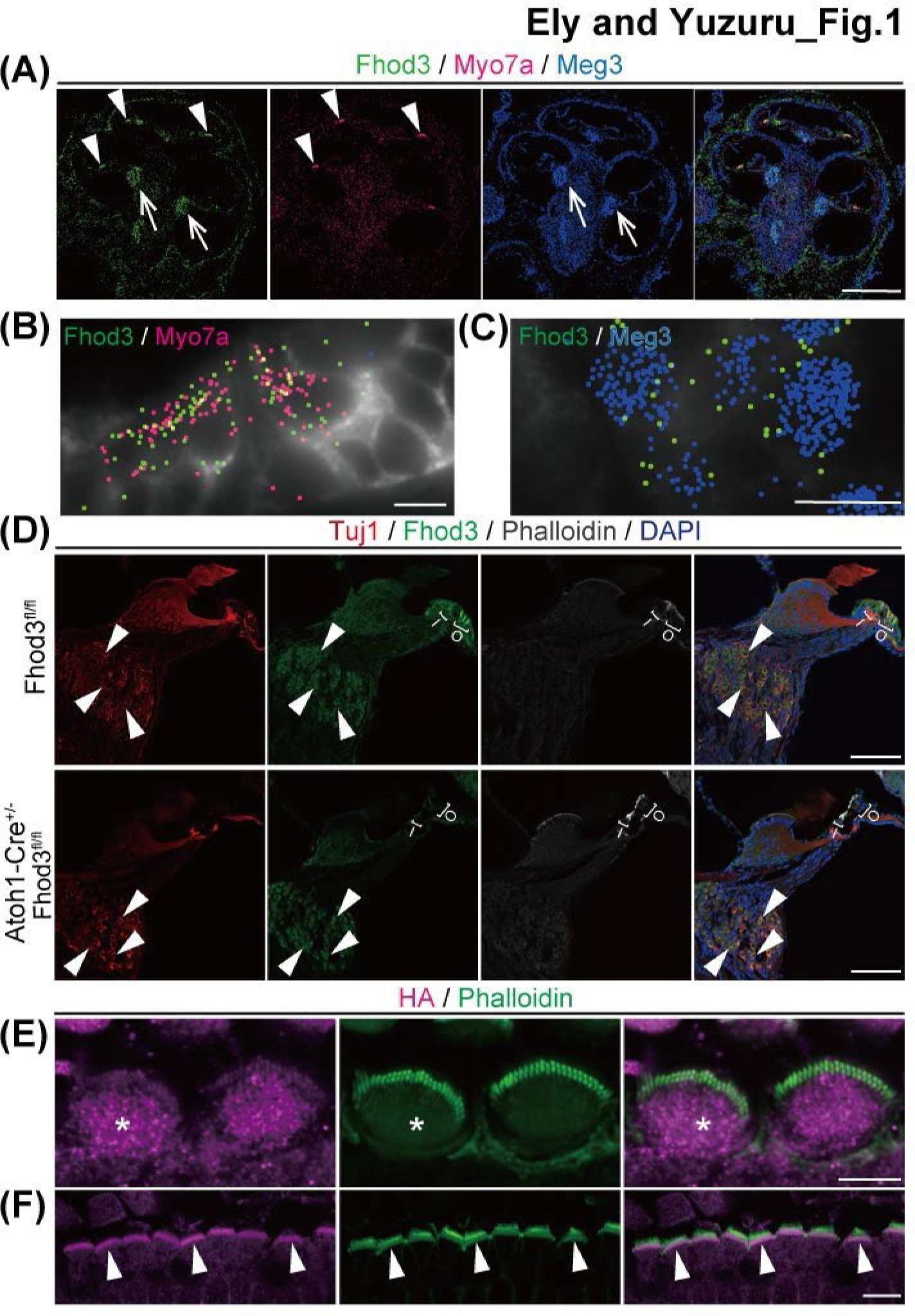
Cellular and Subcellular localization of Fhod3 in the mouse cochlea. Fhod3 mRNA expression in the mouse cochlea at post neonatal day 5(P5) (A-B). **A)** Multiplexed error robust fluorescent in-situ hybridization (MERFISH) image of the mid modiolar section. Fhod3 transcripts (green) expressed in the hair cell (HC) (arrow head) and spiral ganglion cells (SGCs) (arrow). Myo7a is for HC marker (magenta) and Meg3 (dark blue) is for SGN marker. Scale bar; 500µm. **B-C)** The high magnification image of organ of Corti (B) and spiral ganglion cells (C) of Fig.A. Fhod3 expression was observed in both HCs and SGCs. Cell boundaries were stained with cell boundary staining kit (Vizgen) and depicted by grey color. Scale bar; 25µm. **D)** Cryosections of cochleae of wild type (WT) and Fhod3 knock-out (*Atoh1-Cre^+/-^; Fhod3^fl/fl^*) (KO) mice at P14 were stained by Tuj1, Fhod3, phalloidin antibodies and DAPI. Fhod3 expressions were observed in both HCs ant SGCs. Note that Fhod3 signals were diminished in the HC in the KO mouse. Scale bar; 100µm. **E-F)** Outer HCs images at P2 in Fhod3-3xHA knock-in mouse (*Fhod3^HA/+^*). Cochleae were dissected and stained with anti-HA (magenta) and phalloidin (green) antibodies. Fhod3 signals were mainly observed in cuticular plate (asterisk) (E). Lateral view of outer hair cells and arrow heads indicate HA-positive cuticular plate (arrow head) (F). Scale bar: (E) 5µm, (F) 10µm.

### 2.3 Generation of a transgenic mouse model overexpressing Fhod3

In our meta-analysis GWAS, the largest sample set utilized backcross data with C57/Bl6 (B6) and DBA/2J (D2) strains to map *Ahl8* (Ohmen et al. 2014). Considering the hearing effect observed in our meta-analysis, we aimed to investigate the potential differential expression of *Fhod3* as a plausible mechanism. Through quantitative PCR (qPCR) analysis, we found significantly higher levels of Fhod3 mRNA in D2 cochleae, a model of early-onset progressive hearing loss (B6 vs D2; 1.07 ± 0.088 vs 2.00 ± 0.643, *p\*\**=0.0079 by Mann-Whitney U test) (Fig. 2A). These findings led us to hypothesize that *Fhod3* expression variation might contribute to hearing loss. To further explore this hypothesis, we sequenced the *Fhod3* clone obtained from B6 cochleae and identified three missense mutations (A1888C/rs29670988, C2232T/rs13483257, and G2730A/rs30077504) as well as four synonymous polymorphisms in the D2 allele (Fig.2B). Although the coding variations in the D2 allele were not predicted to be deleterious by PolyPhen and SIFT software, we postulated that the hearing loss phenotype could still be associated with this region, possibly due to *Fhod3* overexpression in D2. To investigate this possibility, we generated a transgenic mouse model by introducing the full-length Fhod3 cloned from B6 cochleae between loxP-flanked stop cassettes and IRES-GFP, allowing tissue-specific Fhod3 overexpression in vivo (Fig.2C). Two transgenic mouse lines were obtained and subsequently crossed with a B6 Pax2-Cre driver line (Ohyama and Groves 2004) to induce Fhod3 overexpression in the developing inner ear epithelium. The resulting offspring carrying both the *Fhod3* allele and *Pax2-Cre* genes (*Pax2-Cre^+/-^; Fhod3^Tg/+^*) exhibited normal survival and did not display any gross morphological phenotype. Through qPCR and immunofluorescence, we observed elevated levels of Fhod3 mRNA (WT vs TG; 1.00 ± 0.311 vs 5.84 ± 0.821, *p\*\**=0.0095 by Mann-Whitney U test) (Fig. 2D) and of FHOD3 protein expression in the mutant cochlea (Fig. 2E).

**Fig.2:**
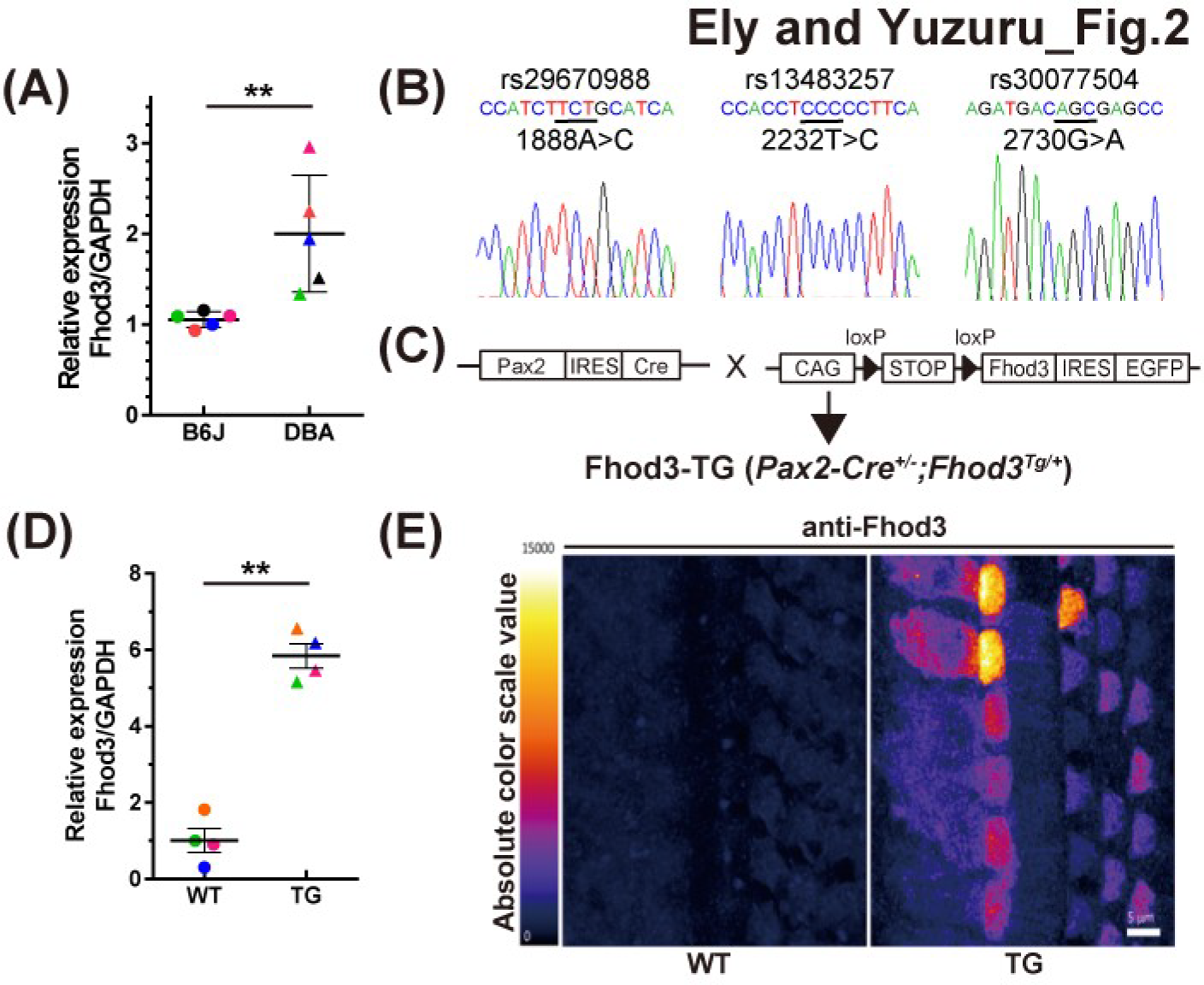
Higher expression and nonsense mutations of Fhod3 in the DBA/2J mice and generation of Fhod3 transgenic mouse. **(A)** Comparison of Fhod3 mRNA expression between C57/BL6 (B6) and DBA/2J (D2) mice. Cochlear mRNA was extracted from P2 mice, and Fhod3 and GAPDH were amplified using gene-specific primers. The relative expression level of Fhod3 was calculated. (n=5, Mann-Whitney U test, *p*\*\*=0.0079). **(B)** Identification of missense mutations in Fhod3 from D2 mice. A comparison of Fhod3 sequence cloned from B6 cochlea revealed three non-deleterious mutations: 1888A>C, 2232T>C, and 2730G>A. Reference SNP (rs) numbers are indicated, and changed residues are underlined. **(C)** Illustration of the generation of the Fhod3 overexpression mouse model. Fhod3 from B6 cochlea was inserted into the CMV/Beta actin Promoter-loxP-STOP-loxP-MCS-IRES-GFP vector, generating a transgenic mouse overexpressing Fhod3 (TG) downstream of the flanked stop codon, controlled by the CMV promoter. The TG mouse was subsequently intercrossed with the Pax2-Cre mouse line to achieve cochlear-specific overexpression of Fhod3 in vivo. **(D)** Relative expression of Fhod3 in wild-type B6 and TG mice. mRNA was obtained from the mice cochleae at P2, and quantitative PCR was performed. Fhod3 expression was normalized to GAPDH. (n=4, Mann-Whitney U test, *p\*\**=0.0095). **(E)** Immunofluorescence intensity of Fhod3 in wild-type (WT) and TG mice at P2. The glow-scale bar demonstrates the color gradient of the scale. Scale bar; 5µm.

### 2.4 Inner ear specific *Fhod3* overexpression leads to progressive high-frequency hearing loss with outer hair cell dominant HC loss and cuticular plate deterioration

To assess the auditory function of the Fhod3 transgenic mice (*Fhod3^Tg/+^; Pax2-Cre^+/-^*), auditory brainstem responses (ABR) were measured at 6 and 12 weeks postnatally. The mutant mice demonstrated a progressive elevation of ABR thresholds specifically at 24 kHz and 32 kHz frequencies [WT vs TG: (6 weeks) 26.7 ± 1.47 vs 38.3 ± 6.91 dB at 24kHz, p= 0.0097, 42.1 ± 7.72 vs 64.6 ± 3.97 at 32 kHz, p<0.0001; (12 weeks) 25.0 ± 1.01 vs 46.25 ± 5.99 dB at 24 kHz, p<0.0001, 42.9 ± 5.28 vs 75.4 ± 2.13 dB at 32 kHz, p<0.0001] (Fig. 3A). Corresponding to the threshold elevation, histological examination revealed HC loss predominantly in the OHCs within the region associated with high frequencies at 5 months old (Fig. 3B). The OHC survival rate in the TG mice was significantly decreased at the basal turn region compared to WT mice (WT vs TG (%): (IHC)100 vs 100 at apical turn (Api), 98.7± 0.95 vs 97.7±1.51 at middle turn (Mid), 94.7±2.24 vs 85.9±6.90 at basal turn (Base); (OHC) 97.1±1.16 vs 95±2.56 at Api, 96.8±0.36 vs 96.5±1.42 at Mid, 95.6±0.64 vs 75.93±8.98 at base, *p*\*=0.0233 by two-way ANOVA followed by a Tukey post-hoc test) (Fig. 3C). Additionally, a significant reduction in phalloidin intensity at the cuticular plate, a critical structure composed of a dense actin meshwork, was observed in OHCs at the mid-basal turn (WT vs TG: (IHC)1.00 ± 0.175 vs 0.92 ± 0.179; (OHC)1.00 ± 0.131 vs 0.70 ± 0.170, *p\*\*\**<0.001, by Mann-Whitney U test). (Fig. 3D and E). These findings strongly suggest that increased Fhod3 expression reduces HC integrity, leading to progressive hearing loss through HC loss.

**Fig.3.**
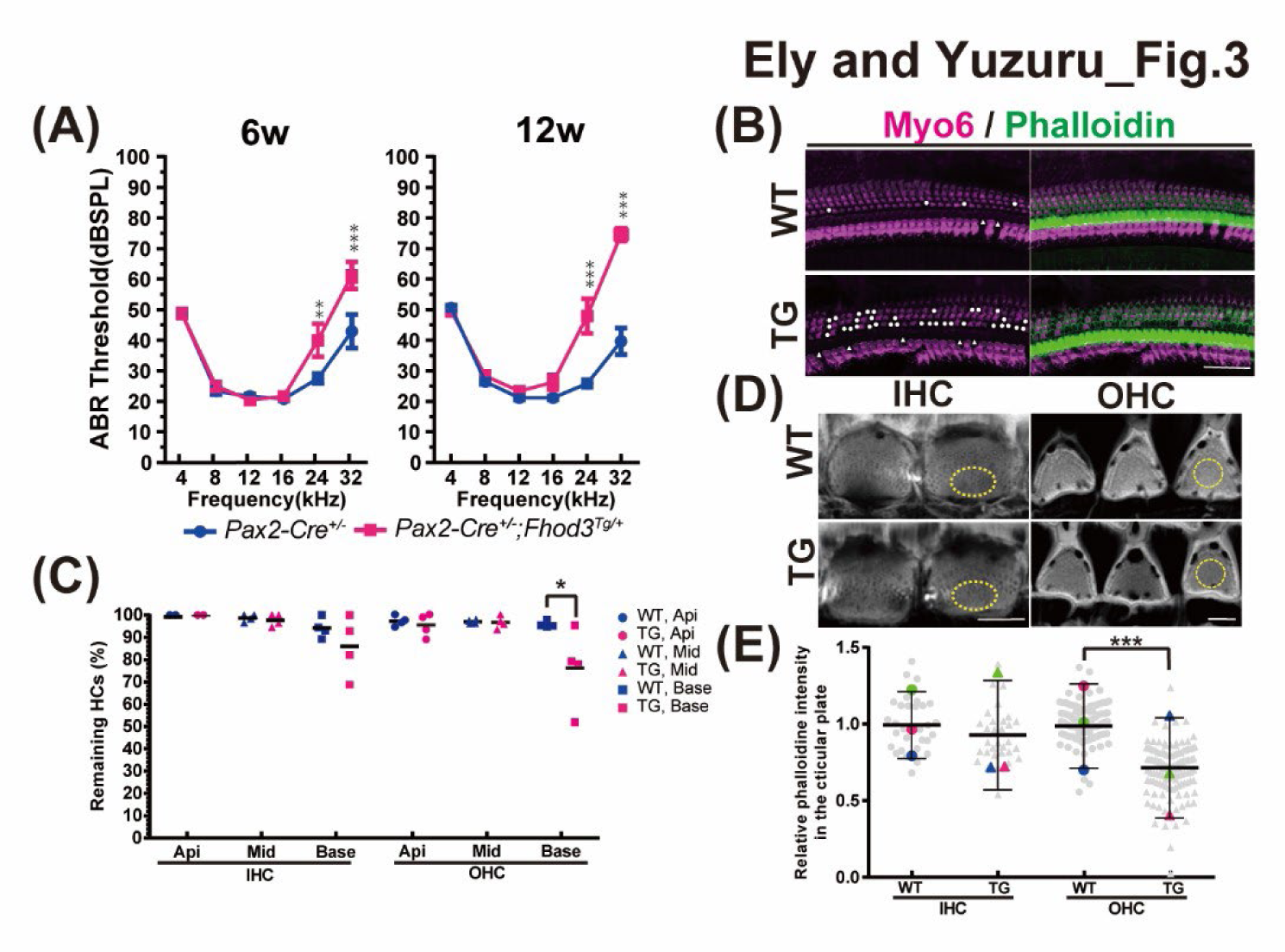
Auditory phenotype and immunohistochemical analysis of Fhod3-TG mice. **(A)** Auditory brainstem response (ABR) thresholds with pure-tone bursts at various frequencies in WT (*Pax2-Cre^-/-^; Fhod3^Tg/+^*) and TG (*Pax2-Cre^+/-^; Fhod3^Tg/+^*) mice at 6 weeks and 12 weeks of age. Significant differences were observed at 24 kHz and 32 kHz, and the thresholds elevation at both frequencies are progressive. Statistical analysis performed using two-way ANOVA followed by a Tukey post-hoc test. (n=12 for each group, *p\*\**=0.0097 at 24 kHz (6 weeks), *p\*\*\**<0.0001 at 32 kHz (6 weeks), *p\*\*\**<0.0001 at 24 kHz (12 weeks), *p\*\*\**<0.0001 at 32 kHz (12 weeks)). **(B)** Representative confocal images of the basal turn cochlea at 5 months old, showing immunostaining with Myosin6 and phalloidin. White dots indicate hair cell losses. Scale bar; 20µm. **(C)** Survival hair cell rate in WT and TG mice at 5 months old. The number of surviving Myo6-positive inner and outer hair cells (IHCs and OHCs) was counted, and the survival rate was calculated in the apical (Api), middle (Mid) and basal (Base) turn cochleae. The survival rate of OHC in TG mice at the basal turn cochlea was significantly decreased compared to the WT control (WT vs TG in the basal OHC (%): 95.6±0.64 vs 75.93±8.98, *p\**=0.023 by Mann-Whitney U test, n=4). **(D)** Immunofluorescence images in the cuticular plate of TG mice and the control at 5 months old. Phalloidin staining (grey) was performed and show representative images of IHC and OHC in the mid-basal turn cochlea. Region of interests for the measurement of phalloidin intensities are depicted by yellow color dots. Scale bar; 5µm. **(E)** Statistical analysis of phalloidin staining in the cuticular plate at 5 months old. Each grey dots indicates relative phalloidin intensities of remaining hair cells (total n=86, 91, 273 and 279), and each colored dots demonstrates average of the intensities per experiment (WT vs TG: (IHC)1.00 ± 0.175 vs 0.92 ± 0.179; (OHC)1.00 ± 0.131 vs 0.70 ± 0.170, *p\*\*\**<0.001, by Mann-Whitney U test, n=3 animals).

### 2.5 FHOD3 overexpression deteriorated the third row of stereocilia of the hair cells in the base of the cochlea

To investigate the impact of Fhod3 overexpression on stereocilia, we conducted an ultrastructural analysis using scanning electron microscopy on transgenic mice (TG) (*Fhod3^Tg/+^; Pax2-Cre^+/-^*) in comparison to wild-type mice (WT) (*Fhod3^Tg/+^; Pax2-Cre^-/-^*) at 7 weeks of age. Our examination revealed a significant decrease in the number of stereocilia in the shortest (third) row of both inner and outer hair cells within the basal regions of the cochlea in the TG mice (Fig. 4). These findings, combined with our localization data demonstrating Fhod3 presence in the cuticular plate (Fig. 1E and F), indicate that FHOD3 plays a crucial role in maintaining actin dynamics within the stereocilia of both IHCs and OHCs via the cuticular plate.

**Fig.4.**
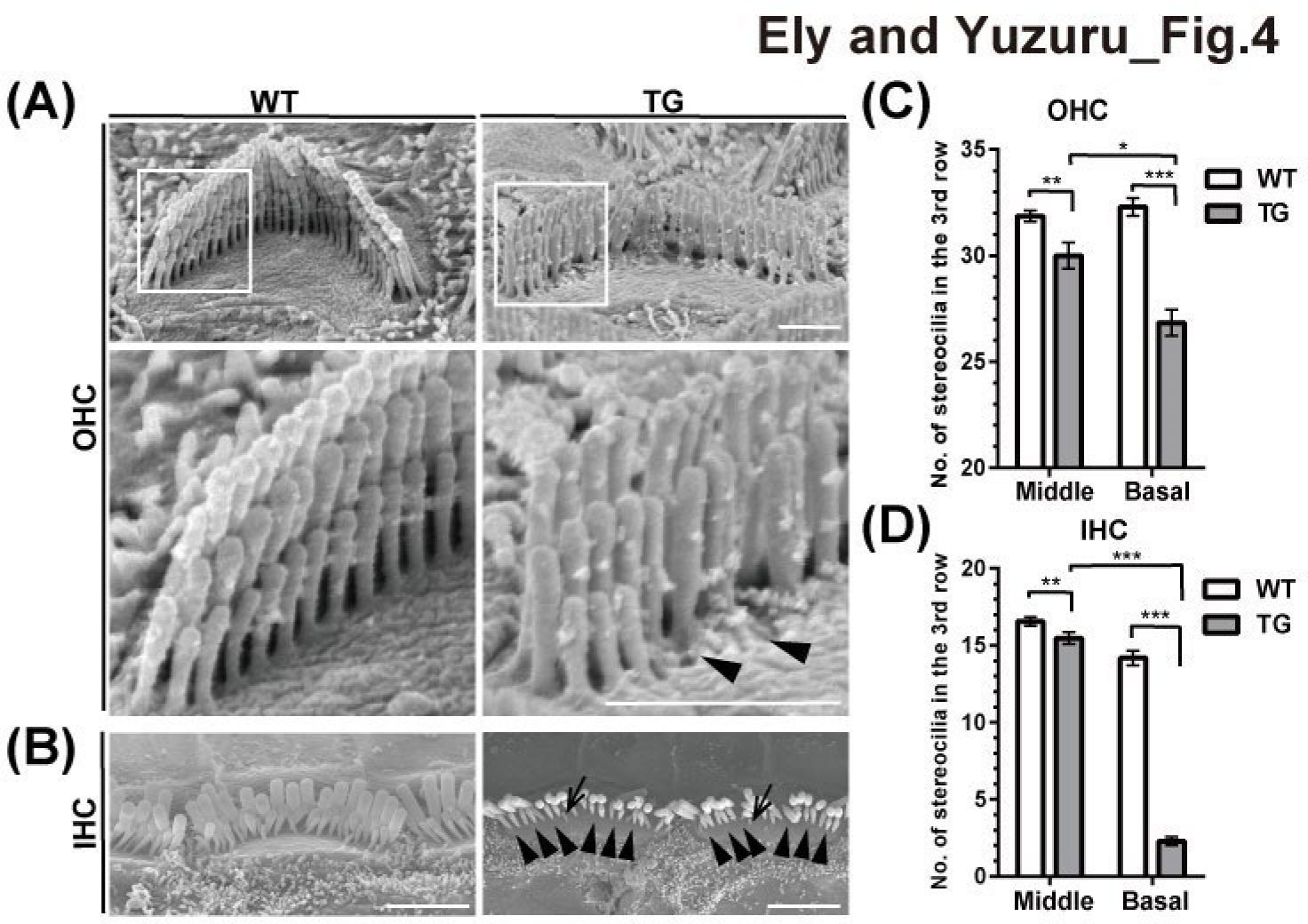
Ultrastructural analysis of Fhod3 transgenic mise. Scanning electron microscopy (SEM) images and the comparison of number of stereocilia in the third row of 7-weeks old age cochleae. **(A) (B)** SEM images of outer hair cell (OHC) (A) and inner hair cell (IHC) (B) hair cell stereocilia in WT (*Pax2-Cre^-/-^; Fhod3^Tg/+^*) and TG (*Pax2-Cre^+/-^; Fhod3^Tg/+^*) mice at basal turn cochleae. White boxed regions are magnified in the lower panel. Arrows indicate missing (arrowhead) and remaining (arrow) third row stereocilia in the TG mice. [Scale bar; (A) 1µm, (B) 2.5µm]. **(C)(D)** Comparison of number of stereocilia in the third row of OHC (C) and IHC (D) at middle and basal turn cochleae. Statistical analysis was performed using Mann-Whitney U test. [(C) n=7 for the middle turn (Mid) and the basal turn (Base) cochlea, *p\**=0.0216 at Mid, *p*\*\*\*<0.0001 at Base; (D) n=9 and 11 for the Mid, n=15 for the Base, *p\*\**=0.0016 at Mid, *p*\*\*\*<0.0001 at Base].

### 2.6 Targeted deletion of *Fhod3* induced progressive hearing loss and hair-cell deterioration

Constitutive knockout of Fhod3 in mice results in prenatal lethality, likely due to developmental defects in the myocardium (Kan-o et al. 2012). Therefore, to gain deeper insights into the specific impact of Fhod3 deficiency on the mouse cochlea, it was crucial to achieve tissue specific inactivation of Fhod3. To accomplish this, we generated mice with a floxed allele of Fhod3 combined with a Atoh1 promoter-driven Cre construct, enabling HC-specific Fhod3 inactivation. Remarkably, these mutant mice exhibited a physiological phenotype similar to that of Fhod3 overexpression mutants, characterized by elevated auditory brainstem response (ABR) thresholds specifically at the highest frequencies of 24 kHz and 32 kHz [WT vs KO: (6 weeks) 34.25 ± 4.12 vs 36.2 ± 2.83 dB at 24kHz, 36.3 ± 5.03 vs 50.1 ± 6.35 at 32 kHz, *p\*\**=0.0096; (12 weeks) 32.3 ± 1.20 vs 52.9 ± 5.80 dB at 24 kHz, *p\*\**=0.0031, 39.0 ± 4.51 vs 73.5 ± 4.55 dB at 32 kHz, *p*\*\*\*<0.0001; (18 weeks) 34.9 ± 2.93 vs 53.8 ± 3.06 dB at 24kHz, *p\**= 0.0187, 51.4 ± 4.91 vs 80.8 ± 3.74 at 32 kHz, *p*\*\*\*<0.0001] (Fig. 5A). Importantly, the observed high frequency hearing loss progressed over time and heterozygous (Het) mice also showed threshold elevation in the high frequency. [Het: (6 weeks) 41.8 ± 6.53 dB at 24kHz, 54.3 ± 5.43 at 32 kHz, *p*\*=0.0179 (vs WT); (12 weeks) 47.0 ± 9.42 dB at 24 kHz, 72.4 ± 6.29 dB at 32 kHz, *p\*\*\**<0.0001 (vs WT); (18 weeks) 67.3 ± 13.6 dB at 24kHz, *p\*\*\**= 0.0002 (vs WT), 86.0 ± 4.86 at 32 kHz, *p\*\*\**<0.0001(vs WT)]. Additionally, we observed a significant increase in hair cell loss in the basal turn of the cochlea at 3 months of age in KO mice (Fig. 5B), along with a notable reduction in phalloidin intensity in IHCs and OHCs at the cuticular plate in the mid-basal turn of the cochlea (WT vs KO: (IHC)1.00 ± 0.037 vs 0.96 ± 0.005, *p\**=0.035; (OHC)1.00 ± 0.023 vs 0.67 ± 0.017, *p\*\*\**<0.001 by Mann-Whitney U test) (Fig. 5C), consistent with our findings in the transgenic overexpression mice (Fig. 3E). Scanning electron microscopy analysis similarly revealed fused, shortened, and missing stereocilia in the mid-base of the cochlea, corresponding to the tonotopic region associated with hearing loss (Fig. 5E). These results indicate that alterations in Fhod3 expression levels, whether elevated or reduced, disrupt the stoichiometry of actin polymerization at the cuticular plate, primarily affecting the HCs of the third row stereocilia in the mid-basal and basal regions of the cochlea.

**Fig.5.**
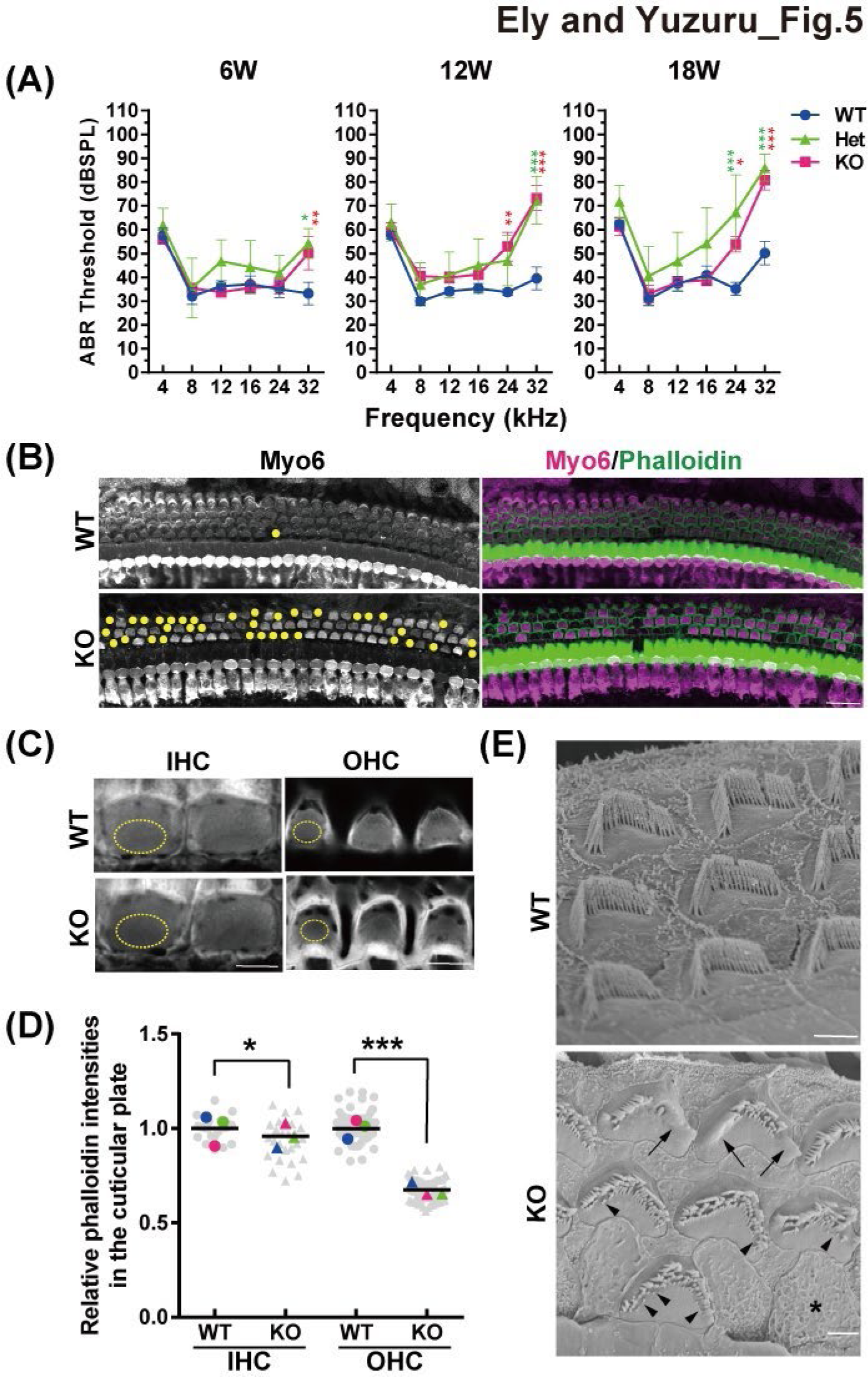
Phenotypic analysis of conditional knock-out mouse of Fhod3. **(A)** ABR measurements of wild type (WT) (*Math1-Cre^-/-^; Fhod3^fl/fl^*) (n=16), heterozygous (Het) (*Math1-Cre^+/-^; Fhod3^fl/+^*) (n=4), knock-out (KO) (*Math1-Cre^+/-^; Fhod3^fl/fl^*) (n=9) mice at 6, 12 and 18 weeks of age. Two-way ANOVA was used to detect significant differences, followed by a Tukey post-hoc test. *p*\*<0.05, *p*\*\*<0.01, *p*\*\*\*<0.001. **(B)** Immunohistochemical analysis was conducted on WT and KO mice cochleae from the basal turn cochlea at 3 months of age. Cochleae were harvested, fixed and stained with Myo6 and phalloidin. Heir cell losses were represented by yellow dots. Scale bar;20µm. **(C)** Representative images of cuticular plate with phalloidin staining in WT and KO mice in the mid-basal turn cochlea at 3-months old. Region of interests for the following phalloidin staining analysis (D) are depicted (yellow dots). Scale bar; 5µm. **(D)** Statistical analysis of phalloidin staining in the cuticular plate at 3-months old. Each grey dots indicates relative phalloidin intensities of survival hair cells (total n=86, 91, 273 and 279), and each colored dots demonstrates average of the intensities per experiment (*p\*\*\**<0.001 by Mann-Whitney U test, n=3 animals). **(E)** SEM images of OHC of WT and KO in the mid-basal-turn cochlea at 3-months old. Abnormal fused stereocilia (arrow), short and missing stereocilia in the third row (arrow head) and complete loss of stereocilia (asterisk) in KO mice are indicated. Scale bar; 2µm.

## 3. Discussion

Here we provide compelling evidence for the crucial role of Fhod3, identified through a meta-analysis of GWAS for age-related hearing loss (ARHL), in maintaining auditory function. Using genetic and molecular approaches, we demonstrated that altered expression of Fhod3 leads to progressive high-frequency dominant hearing loss in two different mouse models: *Pax2-Cre^+/-^; Fhod3^Tg/+^* and *Atoh1-Cre^+/-^; Fhod3^fl/fl^*. Moreover, we observed the deterioration of the cuticular plate and the shortest row of stereocilia in the mutant mice, indicating that dysregulation of Fhod3 disrupts actin turnover balance and affects the integrity of cochlear HCs. This disruption results in the loss of OHCs and could be a potential mechanism underlying ARHL.

In our previous meta-analysis GWAS, which mainly involved C57BL/6J (B6) and DBA/2J (D2) F1 intercross data, we identified five genome-wide significant loci associated with ARHL in mice (Ohmen et al. 2014). In this study, we focused on investigating the potential involvement of Fhod3 in high-frequency and progressive hearing loss, despite the presence of several other candidate genes within the 1Mb interval of the haplotype block. Although we did not find deleterious Fhod3 polymorphisms in the D2 strain (Fig.2B), quantitative PCR revealed significantly higher inner ear expression of Fhod3 in D2 compared to B6 (Fig.2A). Furthermore, Fhod3 mRNA was detected in the human inner ear transcriptome (Schrauwen et al. 2016), supporting its role in human hearing. We subsequently demonstrated that overexpression of Fhod3 in the inner ear leads to high-frequency hearing loss in transgenic mice, accompanied by sparse actin meshwork in the cuticular plate and loss of the shortest row of stereocilia in specific cochlear turn. Notably, mice with P264L gamma actin mutations exhibited a similar phenotype of stereocilia loss (Drummond et al. 2012). While mutations in Fascin Actin-Bundling Protein 2 (Fscn2) were implicated in mid-high frequency hearing loss in D2 mice, they did not account for the complete D2 phenotype, including higher frequency loss and disruption of stereocilia third row in OHCs (Shin et al. 2010). These observations collectively indicate that Fhod3 plays a pivotal role in high-frequency auditory function.

Other formin proteins have been associated with hereditary hearing loss, suggesting that altered expression of Fhod3 in our mutant mouse models may contribute to the underlying molecular mechanisms of hearing loss. Mutations in two formin-related hearing loss genes, DIAPH1 and DIAPH3, are predicted to result in a gain-of-function mechanism (Schoen et al. 2010; Ueyama et al. 2016). In line with this, our Fhod3 overexpression mouse model exhibited progressive hearing loss and dominant loss of OHCs in the basal turn region, resembling the DIAPH1 transgenic mice model (Ueyama et al. 2016; Ninoyu et al. 2020). Considering the subcellular localization of Fhod3 primarily in the cuticular plate, and similar phenotypes between loss and overexpression of Fhod3 leading to defective stereocilia bundle structures, we speculate that Fhod3 expression must be tightly modulated to preserve normal cuticular plate and stereocilia structure. The high frequency range of hearing is particularly susceptible to damage due to its greater mechanical demands but may perhaps also be more generally susceptible to accumulated age-related damage due to higher metabolic requirements (Someya et al. 2010). Significantly, targeted deletion of Fhod3 in conditional knockout mice resulted in a similar phenotype, characterized by elevated ABR thresholds and stereocilia deterioration in the mid-base and base of the cochlea. Even heterozygous mice exhibited progressive hearing loss in the high-frequency range, indicating a dose-dependent effect of Fhod3 expression. Indeed, a Fhod3 variant with a missense mutation at a conserved residue in the FH2 domain (Tyr1249A) causes dilated cardiomyopathy by impairing the ability to induce actin dynamics-dependent activation of the transcription factor serum response factor (SRF) (Arimura et al. 2013). These findings are consistent with a critical role for Fhod3 in maintaining actin stoichiometry in the cuticular plate and stereocilia.

Overall, our study highlights the importance of Fhod3 in regulating actin dynamics and its critical role in maintaining the structural integrity of HCs in the cochlea. These findings provide valuable insights into the pathogenesis of ARHL and open up potential avenues for therapeutic interventions. Further investigations into the molecular mechanisms underlying Fhod3-mediated actin polymerization and its interaction with other proteins involved in hearing loss will deepen our understanding of the disease and pave the way for targeted therapeutic strategies.

## 4 Material and Methods

### 4.1 Animals

All animal protocols were approved (s17178) by the Institutional Care and Use Committee (IACUC) at University of California San Diego (UC San Diego). C57BL/6J and CBA/J mice used in this study were purchased from Jackson Laboratory (ME, USA).

### 4.2 Establish Fhod3 over-expression mouse model

We generated a transgenic (TG) mouse model of Fhod3 expressing CAG-loxP-STOP-loxP(LSL)mouse Fhod3-IRES-GFP. Cochlear RNA was extracted from C57BL/6J mouse with Micro Scale RNA isolation Kit (Ambion) according to the manufacturer’s instructions, followed by cDNA synthesis using ProtoScript® Taq RT-PCR Kit [New England Biolabs (NEB), Ipswich, MA, USA].

*Fhod3* cDNA was amplified with PrimeSTAR® GXL DNA Polymerase (R050B; Takara Bio, San Jose, CA, USA) using a pair of primers (5’-CACCATGGCCACGCTGGCTTGTCGCGTGCAG3’ for the forward and 5’-GAGCATGCTCACAGTTGCAGTTCAGATGTG −3’ for the reverse primer), and subcloned into the pENTR™/D-TOPO™ vector (K2400-20; Thermo Fischer Scientific, Waltham, MA, USA). For the general donor vector, a pair primer (5’-TGCAGGATCCATCGATCACCATGGCCACGCTGGCTTGTCGCGTGCAG-3’ for the forward and 5’-CGAGCTCGTCGACGATTCACAGTTGCAGTTCAGATGTG-3’ for the reverse primer) was utilized for amplifying *Fhod3* cDNA from the plasmid and this 4.7kb fragment was subsequently cloned into a pGreen-RAGE vector cut with EcoR-V, which includes a CMV/Beta actin Promoter-LSL-MCS-IRES-GFP, by using Gibson Assembly® Cloning Kit (NEB, USA). After confirming the identity of the plasmid by sequencing, the linearized construct was electroporated into mouse embryonic stem cells (mESCs) with a C57BL/6 background. Genotyping was performed using the following primer set; 5’-TGCAGGATCCATCGATCACC3’ and 5’-AACTTCGACCCAACGAGC-3’. Fhod3-overexpression mice were generated by crossing *Tg(Pax2-cre)1Akg/Mmnc* mice(Ohyama and Groves 2004) with the *Fhod3^Tg/+^*.

### 4.3 Generation of hair-cell specific Fhod3 conditional knock-out mouse

HC specific Fhod3 conditional knockout mice (KO) were generated by crossing B6.Cg-*Tg(Atoh1cre)1Bfri/J* mice (The Jackson Laboratory, stock no. 011104) with Fhod3^fl/fl^ mice (Accession No. [CDB0927K]: http://www2.clst.riken.jp/arg/mutant%20mice%20list.html) which was generated as previously described (Ushijima T et al. 2018). The Fhod3^fl/fl^ mice contain loxP sites flanking exon 18 of the *Fhod3* gene.

### 4.4 Generation of Fhod3-HA knock-in mice

Using CRISPR/Cas9 editing we generated Fhod3-HA fusion knock-in mice expressing three copies of the human influenza hemagglutinin (HA; TACCCGTATGATGTTCCGGATTACGCT) tag sequence integrated at the C-terminus of Fhod3. Geneious Prime software (Biomatters, Auckland, New Zealand) was utilized to identify the following guide RNA (gRNA) sequences: GCAGTTCAGATGTGCCAACC was located on the last coding exon (exon 28) and GCTGAGCAGAGGACCAGATC was located on the Fhod3 3’ untranslated region (3’UTR). For homology-directed repair (HDR), a 1,879 base-pair Fhod3 donor fragment was amplified from mouse genomic DNA by using the following primers (5’-GCCCATAGGAGGGCCTTTACATTTAGTTTTG-3’and 5’-CAGGTGCCTAGAAAGTGACTTAGATTTCCAGG-3’), and cloned into a custom mini-circle compatible hybrid donor plasmid consisting of two U6 promoters (one for each guide) and a docking site for the homology arms and HA tag as previously described (Jurlina et al. 2022). Integration of the amplified mouse genomic DNA into the donor plasmid and integration of the 3x HA tag before the stop codon were each carried out by Gibson assembly (NEB, USA). HDR into C57BL/6J background mouse embryos was carried out by mixing the plasmid donor, guide RNAs [Integrated DNA Technologies (IDT), IA, USA] and Cas9 protein (Alt-R® S.p. HiFi Cas9 Nuclease V3; IDT, USA) together and microinjecting into mouse embryos with. For mouse genotyping, following a primer set was used; 5’-GCTTTCCACATAGCACCTACA-3’ and 5’-CAGATGTTCAGGTCAAGTCTCC-3’.

### 4.5 Immunofluorescence staining

Mice were anesthetized using an intraperitoneal injection of an anesthetic cocktail (ketamine 92.5mg/kg, xylazine 10 mg/kg, and acepromazine 1.85 mg/kg) and fixed via transcardial perfusion with 4% paraformaldehyde (PFA) in 0.1 M phosphate-buffered saline (PBS) (pH 7.4). The cochleae were harvested and post-fixed overnight at 4 ℃ in 4% PFA in 0.1 M PBS (pH 7.4). After fixation, the cochleae were dissected, and surface preparation was performed. The dissected tissues were permeabilized in PBS with 0.3 % PBST (PBS with 0.3% Triton-X) for 30 min at room temperature, followed by blocking with 10 % normal goat serum (NGS) in 0.03 % PBST for 1 h at room temperature with agitation. Primary antibodies (see below) were applied to the samples in 0.03% PBST with 3% NGS and incubated overnight at 4 ℃. After washing three times with 0.03% PBST for 10 min each, the tissues were incubated with secondary antibodies in 0.03% PBST with 3% NGS for 1 h at room temperature with agitation. The stained tissue was briefly washed with 0.03% PBST and mounted in ProLong Antifade (Thermo Fisher Scientific, Cat#P36970, Carlsbad, CA, USA) with a coverslip. Imaging was performed using a confocal microscope with airy-scan processing (LSM880; Carl Zeiss, Jena, Germany).

For cryo-sectioning, the fixed cochleae were decalcified in 0.12 M ethylenediaminetetraacetic acid (EDTA) at 4℃ for 48 h, followed by cry protection in 30% sucrose at 4℃ overnight. The samples were then embedded in O.C.T. compound (Sakura Finetek, CA, USA) and cryo-sectioned into 10 µm thick slices using a cryostat (CM1860; Leica Biosystems, Nussloch, Germany), The sections were mounted on glass slides, and immunofluorescence staining was performed on the glass slide using a hydrophobic barrier pen (IHC World, Cat# SPM0928, MD, USA), as described above.

### 4.6 Antibodies

The following antibodies were used (polyclonal unless indicated): Fhod3 (Thermo Fisher Scientific, Cat# PA5-23313, 1:100); Myo6 (Proteus Bioscience, Cat# 25-6791, 1:250); anti-HA tag (Abcam, Cat# ab9110, 1:100); TuJ-1 monoclonal (R&D systems, Cat# MAB1195, 1:500); Alexa Fluor 568 conjugated phalloidin (Thermo Fischer Scientific, A12380, 1:250); 4’6369 diamidino-2-phenylindole (DAPI) (Novus Biologicals, Cat# NBP2-31156, 1:50,000); Alexa Fluor 47-conjugated anti-mouse IgG2a (Thermo Fischer Scientific, Cat# A-21241, 1:500); Alexa Fluor 88-conjugated anti-rabbit IgG (Themo Fischer Scientific, Cat# A-11008, 1:500).

### 4.7 Multiplexed error-robust fluorescence *in situ* hybridization (MERFISH)

Samples were prepared following the manufacturer’s instructions (Vizgen, Cambridge, MA, USA). Briefly, cochleae from postnatal day 5 (P5) C57Bl/6J mice were fixed and cryo-sectioned into 10 µm thick slices as described above. RNase inhibitor (NEB, Cat# M0314L, USA) was used to preserve RNA integrity during the dehydration step. The cryosections were placed onto MERSCOPE slide glass (Vizgen, Cat# 20400001, MA, USA) and permeabilized in 70% ethanol at 4 ℃ for 24 hours. Tissues were then incubated with blocking solution (Vizgen, Cat# 20300012, USA) containing cell-boundary staining antibodies (Vizgen, Cat# 20300010 and 20300011, USA) for 1 hour at room temperature. Subsequently, the stained samples were incubated with a customized 140 gene panel mix (Vizgen, Cat# 20300006, USA) including Fhod3 and Myo7a probes for 36 hours at 37 ℃ in a humidified cell culture incubator. After post-encoding hybridization samples were washed with formamide wash buffer (Vizgen, Cat# 20300002, USA), embedded in a polyacrylamide gel incubated with digestion premix (Vizgen, Cat# 20300005, 386 USA), and incubated with clearing premix (Vizgen, Cat# 20300003) until tissues became transparent. Images were acquired and analyzed using the MERSCOPE system (Vizgen, USA). Cell-boundary staining was employed for cell segmentation parameter determination.

### 4.8 Quantitative real-time PCR

Cochleae were harvested at P2 and frozen in liquid nitrogen. Samples were stored at −80 ℃ until following RNA extraction. RNA was isolated with Micro Scale RNA isolation kit (Ambion, Austin, TX, USA) followed by reverse transcription using cDNA synthesis kit (AMIL1791; Ambion, USA). Quantitative real-time PCR (qPCR) was conducted using TaqMan® probe (Fhod3; Mm00614166_m1) obtained from applied biosystems. For each sample, the qPCR assay was performed in triplicate and included the housekeeping gene, GAPDH, as a reference. The 397 relative transcript quantity was determined using the 2^-ΔΔCt^ method, with GAPDH serving as the 398 internal control (Schmittgen and Livak 2008).

### 4.9 Auditory brainstem response

To assess hearing of Fhod3-TG (*Pax2-Cre^+/-^; Fhod3 ^TG/+^*), Fhod3-het (*Atoh1-Cre^+/-^; Fhod3^fl/+^*), Fhod3-KO (*Atoh1-Cre^+/-^; Fhod3^fl/fl^*) mice and littermate control WT mice, auditory brain stem responses (ABRs) were conducted. Mice were anesthetized with the anesthetic cocktail (ketamine 92.5mg/kg, xylazine 10 mg/kg, and acepromazine 1.85 mg/kg), and placed on a heating pad (HP-4M; Physitemp Instruments Inc., Clifton, NJ) regulated by a temperature controller (TCAT406 2DF; Physitemp Instruments Inc, NJ). Subcutaneous stainless-steel electrodes were positioned at the vertex of the skull and the right pinna, with the ground electrode placed near the base of the tail. Acoustic stimuli were generated using a PXI data acquisition system (PXI-1031; National Instruments, Austin, TX, USA) and amplified by an SA1 speaker driver (Tucker-Davis Technologies, Alachua, FL, USA). Sound stimuli were presented to the right ear via an 8-inch long plastic tube attached to a custom acoustic system (Eaton-Peabody Laboratories).(Valero et al. 2018) ABRs were evoked using 5-msec tone pips with a 30/s presentation rate and a 0.5 msec rise/fall Blackman ramp. The stimuli were delivered at sound pressure levels (SPL) ranging from 20 to 100 dB in 5 dB step increments, at frequencies of 4, 8, 16, 24 and 32 kHz. The responses were filtered using a 0.3–3 kHz band-pass filter, amplified by 10,000 times with a P511 preamplifier (Grass Instruments, Astro-Med, Inc., RI, USA), and averaged over 512 responses at each dB SPL. The hearing threshold was determined as the lowest sound intensity level at which a recognizable or reproducible wave was observed.

### 4.10 Quantification of hair cell survival and cuticular plate analysis

The cochleae were immuno-stained with Myo6 and phalloidin, followed by imaging using a 20X objective on an Airyscan confocal microscope (LSM880; Zeiss, Germany) with a pixel size of 49nm. Viable hair HCs labeled with Myo6 were manually counted at each cochlear turn: apical (corresponding to the 6-8 kHz region), middle (corresponding to the 12-24 kHz region), and basal (corresponding to the 32-42 kHz region). The survival rate was calculated as the ratio of the number of Myo6-positive HCs to the total number of HCs, including depleted HCs. ImageJ software (available at http://rsb.info.nih.gov/ij; developed by Wayne Rasband, National Institutes of Health, Bethesda, MD, USA) was utilized for the analysis.

For the cuticular plate analysis, the average phalloidin intensity at the cuticular plate was measured using ImageJ software. Circular regions of interest (ROIs) were defined at the cuticular plate level adjacent to the stereocilia in outer hair cells (OHC) and inner hair cells (IHC). The same ROIs were applied consistently across experiments. Relative intensity changes were calculated by comparing the phalloidin intensity of the mutants with that of age-matched wildtype (WT) mice.

### 4.11 Scanning electron microscopy

All animals used in this work were handled following the NIH and UCSD Guidelines for Animal Care as mentioned earlier. Mice were euthanized with CO_2_ and decapitated. Inner ears were immediately removed with a fine scissor, transferred to a plastic Petri dish where a small aperture in the bone at the apex region was made with a needle to allow the fixative to flow throughout the whole sensory epithelium. A freshly prepared fixative solution containing 4% formaldehyde [Electron microscopy sciences (EMS)], 2.5% glutaraldehyde (EMS), 100 mM sodium cacodylate buffer (EMS), 2 mM CaCl2 was gently injected into the round window and left for 2 h at room temperature. Then, the cochleae were washed in PBS for 20 min three times, organs of Corti fine dissected under a stereoscope, and transferred to 20-mL glass vials with 100 mM sodium cacodylate buffer. Samples were prepared for SEM imaging by OTOT method where two consecutive baths of buffered 1% osmium tetroxide (EMS) and 1% tannic acid (Sigma-Aldrich, St. Louis, MO, USA) were applied for 1 h, with 10 min washes for three times in between. Samples were dehydrated in a graded series of ethanol 200-proof (Thermo Fischer Scientific, USA) till absolute, critical-point dried (Leica EM CPD300; Leica, Wetzlar, Germany), placed on carbon tape onto aluminum stubs, gold-sputtered with ∼4 nm thick (Leica EM SCD500; Leica, Germany), and observed in a FEG-SEM (Zeiss Sigma VP; Zeiss, Germany), operated at 5 kV.

### 4.12 Statistical analysis

All statistical analyses were performed using GraphPad Prism (Prism 9.2.0.; GraphPad Software Inc., La Jolla, CA, USA). All data were presented as mean±standard deviation or standard error of the mean (SEM). For ABR and survival HCs analysis, two-way analysis of variance (ANOVA) was used to detect significant differences, followed by a Tukey post-hoc test. For others, MannWhitney U test was utilized. All statistical details can be found in figure legends. The results were considered statistically significant at P <0.05 (*), P <0.01 (**) and P <0.001 (***).

## 6. Acknowledgement

This study was supported by grants from National Institutes of Health (R01 DC-018566 to R.A.F.). U.M. is supported by NIDCD R21 (DC-018237), NIDCD R01 (DC021075), Core Grant NCI CCSG (CA014195), Nathan Shock Center for Aging Research at the Salk Institute P30 AG068635, and a CZI Imaging Scientist Award.

## 7. Authors Contribution

R.A.F and U.M. designed the experiments. E.B. and Y.N. contributed equally as first author. E.B. performed animal studies and some immunofluorescence (IF). Y.N. performed MERFISH, IF, HCs and cuticular plate analysis. E.B. and Y.N. performed all statistical analysis. Q.L. and N.O. designed and generated the TG mice line. Y.N. and K.J.W. designed and established Fhod3-HA mouse line. R.T. and H.S. designed and generated Fhod3^fl/fl^ mice line. L.A. performed SEM experiments. Y.N., E.B., U.M. and R.A.F. analyzed data and wrote the manuscript.

## 8. Reference

Anttonen T, Belevich I, Laos M, Herranen A, Jokitalo E, Brakebusch C, Pirvola U. 2017. Cytoskeletal Stability in the Auditory Organ In Vivo: RhoA Is Dispensable for Wound Healing but Essential for Hair Cell Development. eNeuro 4: ENEURO.0149-17.2017.

Arimura T, Takeya R, Ishikawa T, Yamano T, Matsuo A, Tatsumi T, Nomura T, Sumimoto H, Kimura A. 2013. Dilated Cardiomyopathy-Associated *FHOD3* Variant Impairs the Ability to Induce Activation of Transcription Factor Serum Response Factor. Circulation Journal 77: 2990–2996.

Benkafadar N, François F, Affortit C, Casas F, Ceccato J-C, Menardo J, Venail F, Malfroy-Camine B, Puel J-L, Wang J. 2019. ROS-Induced Activation of DNA Damage Responses Drives Senescence-Like State in Postmitotic Cochlear Cells: Implication for Hearing Preservation. Mol Neurobiol 56: 5950–5969.

Breitsprecher D, Goode BL. 2013. Formins at a glance. J Cell Sci 126: 1–7.

Chen KH, Boettiger AN, Moffitt JR, Wang S, Zhuang X. Spatially resolved, highly multiplexed RNA profiling in single cells. Science 348:aaa6090.

Drummond MC, Belyantseva IA, Friderici KH, Friedman TB. 2012. Actin in hair cells and hearing loss. Hearing Research 288: 89–99.

E Van Eyken 1, G Van Camp, E Fransen, V Topsakal, J J Hendrickx, K Demeester, P Van de Heyning, E Mäki-Torkko, S Hannula, M Sorri, et al. 2007. Contribution of the Nacetyltransferase 2 polymorphism NAT2*6A to age-related hearing impairment. J.Med. Genet. 44:570–8.

Flint J, Eskin E. 2012. Genome-wide association studies in mice. Nat Rev Genet 13: 807–817.

Fransen E, Bonneux S, Corneveaux JJ, Schrauwen I, Di Berardino F, White CH, Ohmen JD, Van de Heyning P, Ambrosetti U, Huentelman MJ, et al. 2015. Genome-wide association analysis demonstrates the highly polygenic character of age-related hearing impairment. Eur J Hum Genet 23: 110–115.

Friedman RA, Van Laer L, Huentelman MJ, Sheth SS, Van Eyken E, Corneveaux JJ, Tembe WD, Halperin RF, Thorburn AQ, Thys S, et al. 2009. GRM7 variants confer susceptibility to age-related hearing impairment. Hum Mol Genet 18: 785–796.

Fujimoto N, Kan-o M, Ushijima T, Kage Y, Tominaga R, Sumimoto H, Takeya R. 2016. Transgenic Expression of the Formin Protein Fhod3 Selectively in the Embryonic Heart: Role of Actin-Binding Activity of Fhod3 and Its Sarcomeric Localization during Myofibrillogenesis. PLoS One 11: e0148472.

Gates GA, Mills JH. 2005. Presbycusis. The Lancet 366: 1111–1120.

Ghazalpour A, Rau CD, Farber CR, Bennett BJ, Orozco LD, van Nas A, Pan C, Allayee H, Beaven SW, Civelek M, et al. 2012. Hybrid mouse diversity panel: a panel of inbred mouse strains suitable for analysis of complex genetic traits. Mamm Genome 23: 680– 692.

Jurlina SL, Jones MK, Agarwal D, De La Toba DV, Kambli N, Su F, Martin HM, Anderson R, Wong RM, Seid J, et al. 2022. A Tet-Inducible CRISPR Platform for High-Fidelity Editing of Human Pluripotent Stem Cells. Genes 13: 2363.

Kan-o M, Takeya R, Abe T, Kitajima N, Nishida M, Tominaga R, Kurose H, Sumimoto H. 2012. Mammalian formin Fhod3 plays an essential role in cardiogenesis by organizing myofibrillogenesis. Biol Open 1: 889–896.

Lavinsky J, Crow AL, Pan C, Wang J, Aaron KA, Ho MK, Li Q, Salehide P, Myint A, MongesHernadez M, et al. 2015. Genome-wide association study identifies nox3 as a critical gene for susceptibility to noise-induced hearing loss. PLoS Genet 11: e1005094.

Li H, Lu M, Zhang H, Wang S, Wang F, Ma X, Liu J, Li X, Yang H, Shen H, et al. 2022. Downregulation of REST in the cochlea contributes to age-related hearing loss via the p53 apoptosis pathway. Cell Death Dis 13: 1–14.

Lynch ED, Lee MK, Morrow JE, Welcsh PL, León PE, King MC. 1997. Nonsyndromic deafness DFNA1 associated with mutation of a human homolog of the Drosophila gene diaphanous. Science 278: 1315–1318.

Markaryan A, Nelson EG, Hinojosa R. 2010. Major arc mitochondrial DNA deletions in cytochrome c oxidase-deficient human cochlear spiral ganglion cells. Acta OtoLaryngologica 130: 780–787.

Menardo J, Tang Y, Ladrech S, Lenoir M, Casas F, Michel C, Bourien J, Ruel J, Rebillard G, Maurice T, et al. 2012. Oxidative Stress, Inflammation, and Autophagic Stress as the Key Mechanisms of Premature Age-Related Hearing Loss in SAMP8 Mouse Cochlea. Antioxidants & Redox Signaling 16: 263–274.

Miyoshi T, Belyantseva IA, Kitajiri S, Miyajima H, Nishio S, Usami S, Kim BJ, Choi BY, Omori K, Shroff H, et al. 2022.Human deafness-associated variants alter the dynamics of key molecules in hair cell stereocilia F-actin cores. Hum Genet 141:363–382

Ninoyu Y, Sakaguchi H, Lin C, Suzuki T, Hirano S, Hisa Y, Saito N, Ueyama T. 2020. The integrity of cochlear hair cells is established and maintained through the localization of Dia1 at apical junctional complexes and stereocilia. Cell Death & Disease 11: 1–15.

Ohmen J, Kang EY, Li X, Joo JW, Hormozdiari F, Zheng QY, Davis RC, Lusis AJ, Eskin E, Friedman RA. 2014. Genome-wide association study for age-related hearing loss (AHL) in the mouse: a meta-analysis. J Assoc Res Otolaryngol 15: 335–352.

Ohyama T, Groves AK. 2004. Generation of Pax2-Cre mice by modification of a Pax2 bacterial artificial chromosome. genesis 38: 195–199.

Rau CD, Parks B, Wang Y, Eskin E, Simecek P, Churchill GA, Lusis AJ. 2015. High-Density Genotypes of Inbred Mouse Strains: Improved Power and Precision of Association Mapping. G3 Genes|Genomes|Genetics 5: 2021–2026.

Rodríguez-de la Rosa L, Lassaletta L, Calvino M, Murillo-Cuesta S, Varela-Nieto I. 2017. The Role of Insulin-Like Growth Factor 1 in the Progression of Age-Related Hearing Loss. Frontiers in Aging Neuroscience 9. https://www.frontiersin.org/articles/10.3389/fnagi.2017.00411 (Accessed June 7, 2023).

Sánchez-Martínez A, Benito-Orejas JI, Tellería-Orriols JJ, Alonso-Ramos MJ. 2017. Autosomal dominant auditory neuropathy and variant DIAPH3 (c.-173C>T). Acta Otorrinolaringol Esp (Engl Ed) 68: 183–185.

Schmittgen TD, Livak KJ. 2008. Analyzing real-time PCR data by the comparative C(T) method. Nat Protoc 3: 1101–1108.

Schoen CJ, Emery SB, Thorne MC, Ammana HR, Śliwerska E, Arnett J, Hortsch M, Hannan F, Burmeister M, Lesperance MM. 2010. Increased activity of Diaphanous homolog 3 (DIAPH3)/diaphanous causes hearing defects in humans with auditory neuropathy and in Drosophila. Proc Natl Acad Sci U S A 107: 13396–13401.

Schrauwen I, Hasin-Brumshtein Y, Corneveaux JJ, Ohmen J, White C, Allen AN, Lusis AJ, Van Camp G, Huentelman MJ, Friedman RA. 2016. A comprehensive catalogue of the coding and non-coding transcripts of the human inner ear. Hear Res 333: 266–274.

Schuknecht HF, Gacek MR. 1993. Cochlear pathology in presbycusis. Ann Otol Rhinol Laryngol 102: 1–16.

Shin J-B, Longo-Guess CM, Gagnon LH, Saylor KW, Dumont RA, Spinelli KJ, Pagana JM, Wilmarth PA, David LL, Gillespie PG, et al. 2010. The R109H Variant of Fascin-2, a Developmentally Regulated Actin Crosslinker in Hair-Cell Stereocilia, Underlies EarlyOnset Hearing Loss of DBA/2J Mice. J Neurosci 30: 9683–9694.

Someya S, Yu W, Hallows WC, Xu J, Vann JM, Leeuwenburgh C, Tanokura M, Denu JM, Prolla TA. 2010. Sirt3 Mediates Reduction of Oxidative Damage and Prevention of AgeRelated Hearing Loss under Caloric Restriction. Cell 143: 802–812.

Sugiura S, Uchida Y, Nakashima T, Ando F, Shimokata H. 2010. The association between gene polymorphisms in uncoupling proteins and hearing impairment in Japanese elderly. Acta Oto-Laryngologica 130: 487–492.

Taniguchi K, Takeya R, Suetsugu S, Kan-o M, Narusawa M, Shiose A, Tominaga R, Sumimoto H. 2009. Mammalian formin fhod3 regulates actin assembly and sarcomere organization in striated muscles. J Biol Chem 284: 29873–29881.

Ueyama T, Sakaguchi H, Nakamura T, Goto A, Morioka S, Shimizu A, Nakao K, Hishikawa Y, Ninoyu Y, Kassai H, et al. 2014. Maintenance of stereocilia and apical junctional complexes by Cdc42 in cochlear hair cells J Cell Sci 127:2040–52

Ueyama T, Ninoyu Y, Nishio S-Y, Miyoshi T, Torii H, Nishimura K, Sugahara K, Sakata H, Thumkeo D, Sakaguchi H, et al. 2016. Constitutive activation of DIA1 (DIAPH1) via Cterminal truncation causes human sensorineural hearing loss. EMBO Mol Med 8: 1310– 1324.

Ushijima T, Fujimoto N, Matsuyama S, Kan-o M, Kiyonari H, Shioi G, Kage Y, Yamasaki S, Takeya R, Sumimoto H. 2018. The actin-organizing formin protein Fhod3 is required for postnatal development and functional maintenance of the adult heart in mice. J Biol Chem 293: 148–162.

Valero MD, Hancock KE, Maison SF, Liberman MC. 2018. Effects of cochlear synaptopathy on middle-ear muscle reflexes in unanesthetized mice. Hearing Research 363: 109–118.

Van Laer L, Huyghe JR, Hannula S, Van Eyken E, Stephan DA, Mäki-Torkko E, Aikio P, Fransen E, Lysholm-Bernacchi A, Sorri M, et al. 2010. A genome-wide association study for age-related hearing impairment in the Saami. Eur J Hum Genet 18: 685–693.

